# Computational analysis of filament polymerization dynamics in cytoskeletal networks

**DOI:** 10.1101/839571

**Authors:** Paulo Caldas, Philipp Radler, Christoph Sommer, Martin Loose

**Affiliations:** Institute for Science and Technology Austria (IST Austria), Klosterneuburg, Austria

## Abstract

The polymerization–depolymerization dynamics of cytoskeletal proteins play essential roles in the self-organization of cytoskeletal structures, in eukaryotic as well as prokaryotic cells. While advances in fluorescence microscopy and *in vitro* reconstitution experiments have helped to study the dynamic properties of these complex systems, methods that allow to collect and analyze large quantitative datasets of the underlying polymer dynamics are still missing. Here, we present a novel image analysis workflow to study polymerization dynamics of active filaments in a non-biased, highly automated manner. Using treadmilling filaments of the bacterial tubulin FtsZ as an example, we demonstrate that our method is able to specifically detect, track and analyze growth and shrinkage of polymers, even in dense networks of filaments. We believe that this automated method can facilitate the analysis of a large variety of dynamic cytoskeletal systems, using standard time-lapse movies obtained from experiments *in vitro* as well as in the living cell. Moreover, we provide scripts implementing this method as supplementary material.

## I. Introduction

Cytoskeletal proteins related to tubulin and actin play important roles for the intracellular organization of eukaryotic as well as prokaryotic cells. Both protein families bind nucleoside triphosphates, either ATP or GTP, which allows them to assemble into filaments, while the hydrolysis of the nucleotide triggers their disassembly. As a consequence, these cytoskeletal filaments show unique properties distinct from equilibrium polymers, which are essential for various complex processes in the living cell, from chromosome segregation to cell motility.

Microtubules as well as some prokaryotic actin-related proteins, show a behavior termed dynamic instability, where one end of the filament shows phases of growth that are stochastically interrupted by rapid depolymerization events (Deng et al., 2017; Garner, Campbell, & Mullins, 2004; Horio & Hotani, 1986; Mitchison & Kirschner, 1984; Sammak & Borisy, 1988). In contrast, filaments formed by actin and by the bacterial tubulin-homolog FtsZ are known for treadmilling behavior (Fujiwara, Takahashi, Tadakuma, Funatsu, & Ishiwata, 2002; Loose & Mitchison, 2014; Wang, 1985; Wegner, 1976). In this case, nucleotide hydrolysis coupled with a conformational change of the monomer makes the two ends of the filament kinetically different such that the polymer unidirectionally grows from one end, while depolymerization dominates at the opposite end (Wagstaff JM Oliva MA, García-Sanchez A, Kureisaite-Ciziene D, Andreu JM, Löwe J. et al., 2017).

To understand the properties of cytoskeletal filaments and their role for the large-scale organization of the cell, it is essential to study their polymerization–depolymerization dynamics and how they are changed by regulatory proteins. This not only requires experimental assays that can visualize filament dynamics, but also reliable, quantitative methods to measure the rates of polymerization as well as depolymerization in a non-biased, high-throughput manner.

Despite recent advances in high-resolution fluorescence imaging in biology, kymographs are still the most commonly used approach to visualize dynamics, in particular of cytoskeletal structures. In kymographs, the intensity along a predefined path is plotted for every image of a time-lapse movie, which results in the profile at the spatial position of an object over time. This approach not only helps to visualize dynamic processes, but also allows for a direct read-out of speed and direction of a moving object by analyzing the slope of a corresponding diagonal line. Due to their ease of use, kymographs have been routinely used to measure growth and shrinkage velocities of dynamic systems, including the treadmilling behavior of membrane-bound filaments of FtsZ (Loose & Mitchison, 2014; Ramirez et al., 2016). However, using kymographs to quantify polymerization dynamics has serious drawbacks. Despite being assisted by image analysis software, kymographs are essentially a hand-tracking technique, which makes the collection of large data sets not only impractical but also subject to user bias.

Recently, a number of computational approaches have been developed aiming for an automated analysis of experimental data. In these cases, automatic tracking algorithms were designed to either follow the position of microtubules or actin filaments in filament gliding assays (Ruhnow, Zwicker, & Diez, 2011), an important tool to study molecular motors, or microtubule polymerization dynamics (Kapoor, Hirst, Hentschel, Preibisch, & Reber, 2019). However, reliable single filament tracking is currently impossible within dense assemblies of homogeneously labelled filaments, such as in the actin cortex, mitotic spindle or membrane-bound cytoskeletal networks of FtsZ.

To overcome these difficulties, different alternative strategies relying on automated data collection and analysis have been developed. For instance, fluorescently labelled microtubule plus-end binding proteins have been used as markers for plus end-growth (Perez, Diamantopoulos, Stalder, & Kreis, 1999). These proteins specifically recognize a structural property of the growing end of the microtubule, where they form a characteristic comet-like fluorescence profile. These comets were first semi-manually tracked and analyzed (Gierke, Kumar, & Wittmann, 2010), but later improved image analysis methods helped to fully computationally detect and track EB1 comets in space and time. This made possible to analyze large populations of microtubule ends in an unbiased manner even in the crowded environment of a mitotic spindle, microtubule asters in cells or cytoplasmic extracts (Applegate et al., 2011; Matov et al., 2010). However, up to date the ability to use filament end binding proteins as a proxy for polymerization dynamics only exists for microtubules and is not available for other cytoskeletal systems. Furthermore, depolymerization velocities are not directly available with this method. A more generally applicable approach was introduced with Fluorescent Speckle Microscopy, which has been proven to be valuable for analyzing *in vivo* dynamics of actin and microtubule sliding in living cells as well as cell extracts (Waterman-Storer, Desai, Chloe Bulinski, & Salmon, 1998). Here, small amounts of fluorescently labelled molecules are added to an endogenous dark background, producing a random distribution of fluorescent spots inside cytoskeletal structures. Intensity changes in these speckles disclose information regarding the turnover and binding constants of the subunits, while their motion can be computationally analyzed to retrieve an intracellular map of polymer dynamics. However, fluorescence speckle microscopy does not provide a direct readout of filament growth and shrinkage. Instead, in the case of actin, polymerization rates are derived from the quantification of retrograde polymer flow, while depolymerization is estimated from the decay of speckle number with time (Watanabe & Mitchison, 2002). Accordingly, in the absence of polymer transport, a quantification of polymerization dynamics is not possible by Speckle Microscopy.

To address these limitations, we sought to develop a new image analysis workflow that would allow us to track polymerization dynamics in a more accurate and automated manner using time-lapse movies of homogeneously labeled filaments. We describe here a computational image analysis strategy to directly quantify the polymerization and depolymerization of treadmilling FtsZ filaments using established experimental data acquisition protocols. Our approach relies on standard time-lapse imaging and open-source software tools, which makes it readily available to study various dynamic processes *in vivo* and *in vitro*.

## II. Approach & Rationale

The bacterial tubulin FtsZ plays a central role in the spatiotemporal organization of the bacterial cell division machinery. It assembles into a ring-like cytoskeletal structure at the center of the cell, (Filho et al., 2016; Loose & Mitchison, 2014; Yang et al., 2017), where treadmilling filaments of FtsZ recruit and distribute the proteins required for cell division. To study the intrinsic polymerization dynamics of FtsZ filaments, we routinely use purified proteins in an *in vitro* system based on supported lipid bilayers (SLBs) combined with TIRF microscopy (Fig. 1A). In this experimental setup, FtsZ and its membrane anchor FtsA form treadmilling filaments on the membrane surface, which further organize into rotating swirls and moving streams of filaments (Fig. 1B) (Loose & Mitchison, 2014). As illustrated in Figure 1C, obtaining the velocity of FtsZ treadmilling from kymographs involves selecting high contrast features, drawing a line along the treadmilling path and extracting the velocity from the slope. Due to the manual nature of all of those steps, this procedure is highly sensitive to measurement errors.

**Fig. 1:**
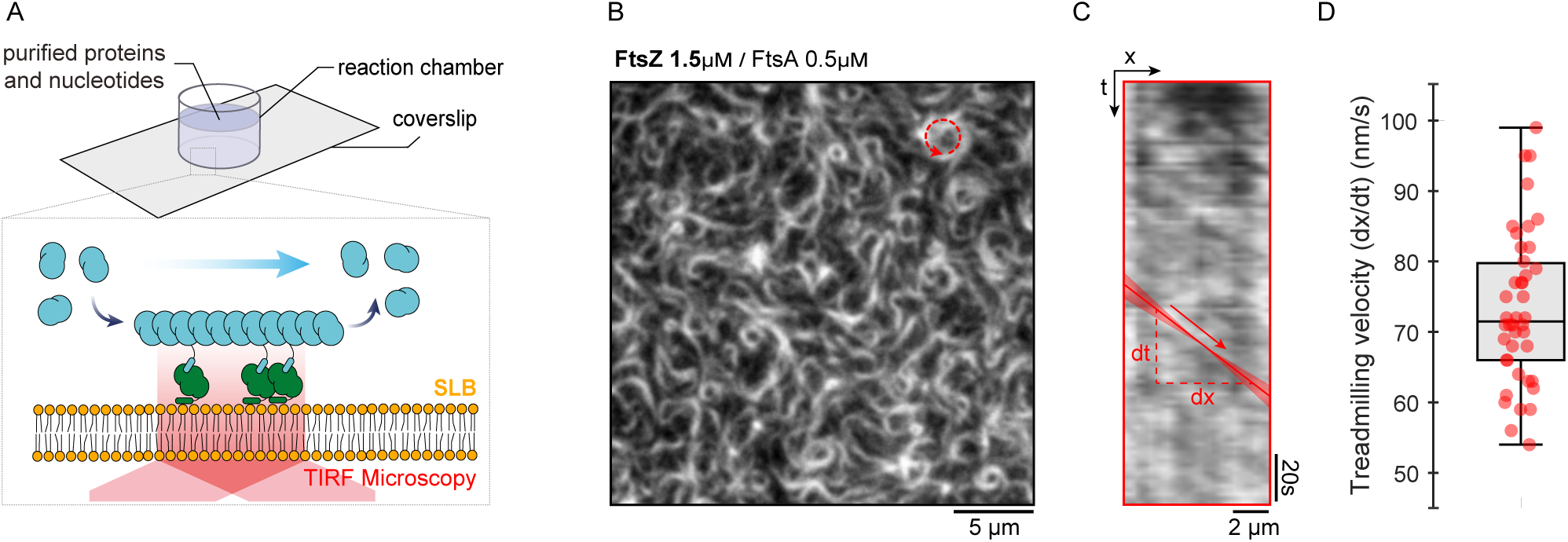
Time-lapse imaging of dynamic FtsZ filaments. **(A)** Illustration of the experimental assay based on a supported lipid bilayer, purified proteins and TIRF microscopy. **(B)** Snapshot of FtsZ (30% Cy5-labelled) pattern emerging from its interaction with FtsA, 15min after the addition of GTP and ATP to the reaction buffer. Scale bars, 5µm. **(C)** Representative kymograph of treadmilling dynamics taken along the contour of a rotating FtsZ ring. The shaded bar illustrates the imprecision associated with manually drawing the diagonal line to estimate velocity (red). Scale bars, x = 2µm, t = 20s.

The workflow of our new protocol to analyze filament polymerization dynamics starts with time-lapse movies of evenly labelled FtsZ filament networks, followed by three computational steps: (i) generation of dynamic fluorescent speckles by image subtraction; (ii) detection and tracking of fluorescent speckles to build treadmilling trajectories and (iii) analysis of trajectories to quantify velocity and directionality of filaments. Importantly, after an initial optimization of parameters, this protocol can be used in batch to access data of thousands of trajectories in a highly automated manner.

### a) Image Acquisition

As for all fluorescence microscopy-based approaches, imaging conditions have to be optimized to avoid excessive photo-bleaching while maintaining a good signal-to-noise ratio. Furthermore, the acquisition frame rate needs to be sufficiently high to follow the dynamic process of interest. Detailed experimental protocols to acquire FtsZ treadmilling dynamics can be found in previous publications of this book series (Baranova & Loose, 2017; Nguyen, Field, Groen, Mitchison, & Loose, 2015). For FtsZ treadmilling on supported bilayers, we identified a rate of 2 seconds per frame as well suited to generate well-defined fluorescent speckles. At this rate, an FtsZ filament is known to grow for 60 - 100 nm, i.e. about the size of a pixel in our TIRF microscope setup that can be followed in time allowing for unambiguous trajectory building.

### b) Generating fluorescent speckles by differential imaging

So far, due to the high intensity background in bundles of homogeneously labelled FtsZ filaments, polymer ends are difficult to identify, which has made the automated detection and analysis of polymerization impossible. To extract dynamic information from our time-lapse movies, we took advantage of a background subtraction method also used for motion detection in computation-aided video surveillance (Singla, 2014). By subtracting the intensity of two consecutive frames, a new image is created where persistent pixels are removed and only short-term intensity changes are kept. Thus, non-moving objects generate small absolute pixel values, while high positive and negative intensity differences correspond to fluorescent material being added or removed at a given position, respectively. When applying this procedure to our image sequences, we generate a new time-lapse movie containing moving fluorescent speckles that correspond to growing and shrinking filament ends within FtsZ bundles. Accordingly, this process allows to visualize and quantify polymerization as well as depolymerization rates (Fig. 2).

**Fig. 2:**
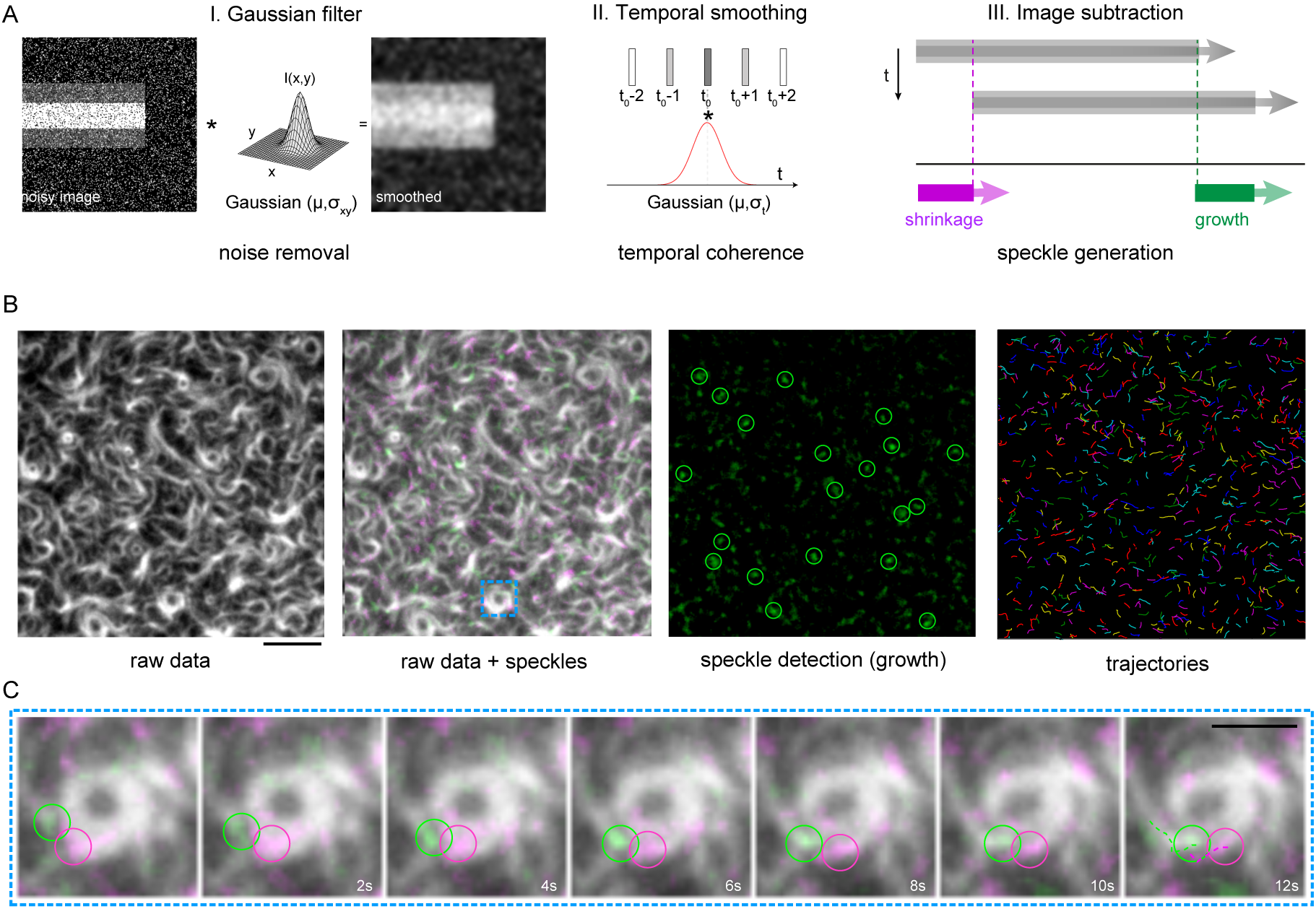
Generating fluorescent speckles by differential imaging. **(A)** Illustration of the Gaussian filtering step in space (I) and time (II) before image subtraction (III). The smoothing in space and time is proportional to σ_xy_ and σ_*t*_, respectively, and μ = 0 represents the mean value of the distribution. **(B)** Representation of differential TIRF images to visualize polymerization (growth) and depolymerization (shrinkage) of FtsZ filaments and automated tracking of treadmilling dynamics with TrackMate (ImageJ). Scale bars, 5µm **(C)** Characteristic detection and tracking of fluorescent speckles inside bundles of FtsZ ring-like structures. Scale bars, 2 µm.

While this process can be easily applied using the ImageJ ImageCalculator plugin, simple image subtraction is susceptible to noise and may generate stretched speckles when the sample acquisition rates are not ideal. Therefore, we incorporated a pre-processing step where we apply a spatiotemporal low-pass filter prior to image subtraction (Fig. 2A). This procedure uses a 3D Gaussian filter where the extent of the smoothing is defined by *σ*_*xy*_ and *σ*_*t*_, representing a convolution in space and time, respectively. The spatial smoothing replaces each pixel value by the Gaussian-weighted average of its neighboring pixels, while the temporal filter replaces each pixel value by the averaged pixel intensity in the previous and subsequent frames. This spatiotemporal smoothing not only effectively removes acquisition noise, but also improves speckle detection and tracking in the next step (Fig. 2B, C).

### c) Fluorescent speckle detection and trajectory building

To quantitatively describe the dynamic behavior of the fluorescent speckles generated in the previous step, we took advantage of particle tracking methods, a common tool to quantitatively analyze the dynamics of moving objects in fluorescent microscopy experiments. These tools are typically implemented either as scripts for programming languages or as plugins for the widely-used software platform ImageJ (Schindelin et al., 2017; Schneider, Rasband, & Eliceiri, 2012) or its distribution FIJI (Schindelin et al., 2012). They usually work by first detecting bright particles in individual images of time-lapse movies and then reconstructing trajectories from the identified spatial positions in time (Fig. 2B). Finally, these trajectories can be further analyzed to retrieve quantitative information about the type of behavior (e.g. directed or diffusive motion), diffusion constant, velocity or lifetime of the particles, as well as the length of trajectories. In our analysis, we chose to use TrackMate for particle detection and tracking (Tinevez et al., 2016), not only because it is an open-source toolbox available for ImageJ, but also because it provides a user-friendly graphical user interface (GUI) with several useful features for data visualization, inspection and export.

### d) Post-tracking analysis

The output of the tracking processes corresponds to the spatiotemporal coordinates of growing and shrinking filaments ends. To quantify their speed and directionality, we can then compute their mean squared displacements (MSD, 〈 ∆*r*^2^ 〉), which is given by the squared displacement as a function of an increasing time lag (*δt*):

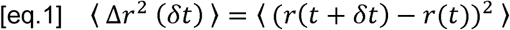

where *r*(*t*) is the position of the particle (*x, y* coordinates) at time *t*, for a given trajectory. MSD curves are easy to interpret and can be mathematically described by diverse models of motion (Qian, Sheetz, & Elson, 1991). Most commonly, the MSD is calculated by averaging all trajectories into one weighted curve, whose shape provides information about the type of motion, e.g. random motion or directional movement. For instance, for randomly moving particles, the MSD curve corresponds to a straight line and can be fitted to:

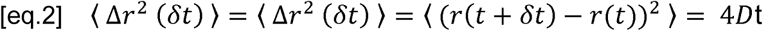

where *D* corresponds to the diffusion constant of the particles. For objects moving directionally, the MSD curve shows a positive curvature and can be fitted to a quadratic equation containing both a diffusion (D) and a constant squared velocity (*v*) term:

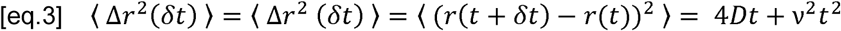

Irregular trajectories containing immobile or confined particles require other models (Qian et al., 1991). In agreement with the directionality of treadmilling, MSD curves obtained from our experiments displayed a positive curvature and were always best fitted to quadratic equations (Fig. 3A).

**Fig. 3:**
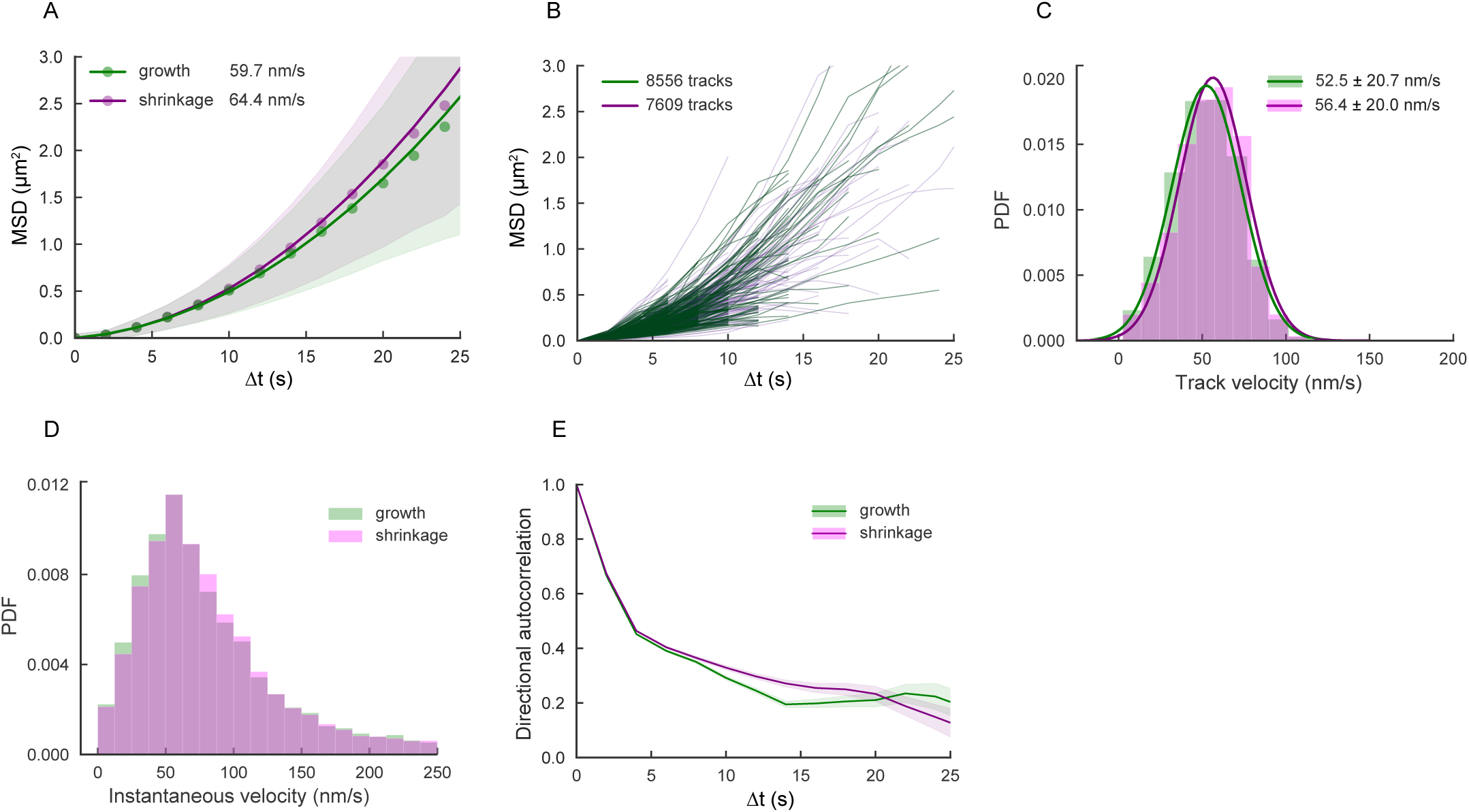
Quantitative analysis of speckles trajectories. **(A)** weighted-mean MSD curve estimated from all trajectories for growth (green dots) and shrinkage (magenta dots) movies. Solid lines represent fitting the data to a quadratic equation (eq.3) to estimate the mean velocity of the speckles. We used 50% of the max track length for fitting and the shade bars correspond to the standard deviation of all the tracks. **(B)** Individual MSD curves for growth (green lines) and shrinkage (magenta lines) movies. **(C)** Distribution of velocity values obtained from fitting eq.3 to 50% of each individual MSD curve in B, for both growth (green) and shrinkage (magenta) movies. Data is fitted to a Gaussian function (solid lines) to estimate mean velocity value. **(D)** Distribution of velocity values obtained from the step size displacements of speckles between consecutive time points. **(E)** Velocity auto-correlation analysis using the definition in eq.4. Correlation function shows positive values even for larger *δt*, for both growth and shrinkage events, characteristic of particles moving directionally.

Alternatively, it is possible to compute MSD curves for each trajectory individually (Fig. 3B) and use the corresponding fitting parameters to calculate a histogram of treadmilling velocities (Fig. 3C). This approach has the advantage of revealing outliers and provides a way to distinguish between different subpopulations in the data.

The mean speed of treadmilling can also be obtained by calculating the histogram of instantaneous displacements of every detected particle between two consecutive frames. For FtsZ treadmilling, this gives rise to a positively skewed normal distribution that peaks at a velocity similar to the one obtained from the MSD fits with a long tail towards faster values (Fig. 3D).

We further corroborated the directionality of the trajectories by computing the directional autocorrelation function (Fig. 3E). The corresponding correlation coefficient (*φ*) is a measure for the local directional persistence of treadmilling trajectory *i* and is obtained by computing the correlation of the angle between two consecutive displacements 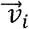 as a function of an increasing time interval (*δt*) (Gorelik & Gautreau, 2014; Qian et al., 1991).

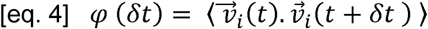

Randomly moving particles typically show completely uncorrelated velocity vectors with *φ* = 0 for all *δt*, while directed moving particles have display highly correlated velocity vectors (*φ* > 0) even for larger *δt*. Accordingly, for our fluorescent speckles we obtained continuously positive correlation values.

## III. Image Analysis using ImageJ and Python

To analyze multiple time-lapse movies of treadmilling filaments in an automated manner, we developed macros based on ImageJ plug-ins and Python. These are easy to use, require no programming knowledge and are available as supplementary material on Github: https://github.com/paulocaldas/Treadmilling-Speed-Analysis.

This software package is organized into three different computational steps:

i. **extraction**: ImageJ macro to automatically generate fluorescent speckles from time-lapse movies;
ii. **tracking**: ImageJ macro that internally uses TrackMate for detection and tracking of fluorescence speckles;
iii. **tracking_analysis**: Python package providing an IPython notebook with detailed analysis of the trajectories generated by TrackMate.

All these scrips can be applied for a single time-lapse movie or for multiple files at once in batch processing mode. This creates a highly time-efficient routine to identify and track thousands of speckles at once. Below, we provide a simple protocol on how to use the macros.

### a) Extraction: Generating fluorescent speckles with ImageJ

To generate speckles from a single time-lapse movie:

1. Open movie of interest in ImageJ (or Fiji).
2. Open macro `extract_growth_shrink.py` and run it.
3. A window pops up to correct/confirm the physical units (i.e. pixel-width and frame interval).
4. A dialog is displayed to set the frame range for the image subtraction – processes the entire movie by default - and the Gaussian smoothing parameters *σ*_*xy*_ and *σ*_*t*_. Generally, the extent of the spatial smoothing is defined by the standard deviation (σ) of two Gaussian functions (σ_x_ and σ_y_). Our protocol applies isotropic smoothing (σ_x_ = σ_y_ = σ_xy_) and should be adjusted according to the size of the object of interest (in pixels). Likewise, the number of frames considered for temporal filtering (σ_t_) depends on the dynamics of the process studied and needs to be optimized for the given frame rate. This parameter is adjusted through trial and error until speckles with a good signal-to-noise ratio are created. For our images we used σ_xy_= 0.5 pixels and σ_t_ = 1.5 frames.
5. The output is a composite movie containing two new channels corresponding to growth (green) and shrinkage (red) together with the raw data (blue). These channels can be split and used for the following analysis step (Fig. 2A).

Once the optimal Gaussian smoothing parameters are defined for a given experimental setup, this process can be applied for several files at once in batch process mode:

1. Open macro `extract_growth_shrink_batch.py` in ImageJ (or Fiji) and run it.
2. A dialog is displayed to select a directory containing the input movies and set the parameters for batch analysis. By default. tif and. tf8 files are processed, if no other file extension is provided. As before, set the frame range, calibrate the physical units, and provide the optimal parameters (*σ*_*xy*_ and *σ*_*t*_) ideally determined beforehand using the macro for a single movie.
3. Gaussian smoothing and image subtraction are then applied for every file in the input directory and two time-lapse movies containing fluorescent speckles (growth and shrinkage) are saved to disk.

### b) Tracking: Building trajectories with TrackMate

When applying this routine for the first time, TrackMate GUI should be used to identify the optimal parameters for detecting, tracking and linking the trajectories of fluorescent speckles. In contrast to single molecule imaging, fluorescent speckles generated by image subtraction can vary a lot in shape and intensity. Therefore, care must be taken to find the right parameters and to discard noise due to random motion. Detailed documentation on how to use TrackMate can be found in (Tinevez et al., 2016) as well as online (https://imagej.net/TrackMate). Nevertheless, we briefly describe here our rationale to find the optimal parameters using TrackMate GUI:

1. Open a differential movie obtained from step a) in ImageJ (or Fiji).
2. Run TrackMate GUI (from Plugins menu)
3. The first panel displays space and time units from the file metadata. They should be accurate as they were calibrated in the previous step when generating the fluorescent speckles. This is important as every subsequent step will be dependent on this calibration and not on the pixel units anymore.
4. Select the Laplacian of Gaussian detector (LoG) for particle detection.
5. Tune the particle’s *estimated diameter* using the ‘preview’ tool, which overlays all detections with circles (magenta) having the set diameter. Note that a high number of low quality spots are erroneously detected by default. They can be easily discarded increasing the *threshold* parameter. Check box for the median filter and sub-pixel localization to improve the quality of detected spots. For speckles obtained from FtsZ treadmilling, TrackMate achieves robust detection using a spot diameter of 0.8 microns and a threshold of 1. Following the detection process, we set an additional filter to keep only particles with a signal-to-noise ratio higher than 1.
6. For trajectory building, we use the Simple Linear Assignment Problem (LAP) tracker algorithm, which requires three parameters: (i) the *max linking distance*, (ii) the *max distance for gap* closing and (iii) the *max frame gap*. The first parameter defines the maximally allowed displacement between two subsequent frames. This value has to be chosen carefully, as too low values result in fragmentation of trajectories with large displacement steps, while too large values can lead to erroneous linking. The other two parameters consider spot disappearance when building trajectories, which can be caused by focus issues during data acquisition. The temporal Gaussian filter applied in our pre-processing steps minimizes any gaps in the treadmilling trajectories. Accordingly, we set *max linking distance* to 0.5 µm and *max distance for gap* closing to 0.
7. After the trajectories are built, we exclude trajectories shorter than six seconds and trajectories with a total displacement smaller than 0.4 µm (half of the particle diameter) to avoid tracking fluorescent speckles not corresponding to treadmilling events.
8. At this point, TrackMate offers a set of interactive tools to examine spots and tracks, which are useful to evaluate the quality of the tracking process, revisit the procedure and adjust some of the parameters. All trajectories are then exported as an XML file (last panel), which contains all the identified treadmilling tracks as spot positions in time.

Once the parameters for a given experimental setup are defined, the TrackMate protocol can be applied to multiple time-lapse movies simultaneously with our ImageJ macro named `track_growth_shrink_batch.py.` This macro provides a GUI to select a directory folder containing growth/shrinkage movies and to define all the parameters described before. TrackMate runs windowless for all files and saves the resulting XML file containing spot coordinates in the same directory. In addition, a TrackMate file suffixed ‘_TM.xml’ is generated, which can be loaded into ImageJ using the “Load TrackMate file” command and allows to revisit the whole analysis process for each file individually.

### c) Tracking analysis: Quantifying speed and directionality

Our routine to analyze speckle trajectories was implemented in Python and can be used from an IPython notebook. It contains two main functions: one to analyze a single XML file and a second one to analyze a folder containing multiple XML files at once. All imported modules located in the adjacent folder can be edited and adapted according to the needs of each user. Our code depends on Python >= 3.6 and we recommend to use the Anaconda python distribution.

To install all the required Python packages for the analysis:

1. Clone/download the Github repository tracking_analysis to a given directory.
2. Open Anaconda prompt in the start menu (command line) and change the directory to tracking_analysis folder by typing:

~~~
cd path_to_directory\tracking_analysis
~~~
3. Install all necessary Python modules locally by writing (note the dot at the end):

~~~
pip install –r requirements.txt –e.
~~~
4. All requirements are automatically resolved.

To use the notebook:

1. Open analyze_tracks.ipynb in Jupyter or IPython notebook.
2. The first module uses a single XML file (TrackMate output) as input, set by the variable “filename” inside the function. This script computes: (i) the weighted-mean MSD curve and estimates velocity by fitting the data to eq.3; (ii) the distribution of velocities from fitting MSD curves individually with eq.3; (iii) the distribution of velocities directly estimated from spot displacement inside each track; (iv) the velocity auto-correlation analysis using the definition on eq.4. All plots are displayed and saved into a single pdf file along with an excel book containing all the data to plot elsewhere if needed.
3. The second module uses a directory as input and runs the exact same process for every XML file inside. As above, all the plots and raw data are saved in a pdf file and excel book, respectively. Note that our protocol assumes directed motion of the speckles and for that reason, MSD curves in our Python script are always fitted to eq.3, as described before. Moreover, as the lag time increases, the number of MSD coefficients available for averaging decreases, producing poor statistics for higher lag times. For this reason, MSD curves are typically fit to less than 50% of the total length of the trajectories. In our notebook, `clip` is an adjustable parameter that defines the % of the trajectory length to be fitted and is set to 50% by default (clip = 0.5), the value used in our analysis. A second optional parameter named `plot_every` can be tuned. This is inversely proportional to the number of individual MSD curves to plot, which can help in dealing with a crowded plot and computation time.

An optional feature allows to run this analysis using the command line interface:

1. Open Anaconda prompt and change directory to tracking_analysis (as above)
2. Run the Python command line interface. e.g for example file with clip = 0.25:

~~~
analyze_tracks_cli example\example_growth_Tracks.xml --clip 0.25
~~~

The final output of our program are five individual graphs showing the instantaneous velocities of growing and shrinking ends, MSD curves of individual trajectories and corresponding histogram, a plot of the weighted average of all MSD curves, as well as the directional autocorrelation. In sum, our macros provide a detailed analysis of filament polymerization dynamics.

## IV. Other applications: Tracking microtubule growth

Our method was initially developed to track treadmilling dynamics of FtsZ filaments, but is applicable to other dynamic filament systems as well. As an example, we chose analyze the growth dynamics of microtubules in Xenopus egg extracts (see Fig. 4). Image subtraction of the original movie gives rise to fluorescent speckles at the growing end of microtubules, which have an appearance reminiscent to EB1 comets. Automated tracking of these speckles shows that the average microtubule growth speed under these conditions was 7.3 ± 2.9 µm/min. This value was obtained within a couple of minutes from the analysis of more than 500 trajectories and is similar to previous reports, where growth was quantified using 50 to 100 kymographs (Thawani, Kadzik, & Petry, 2018). Depolymerization events were not detected using this method.

**Fig. 4:**
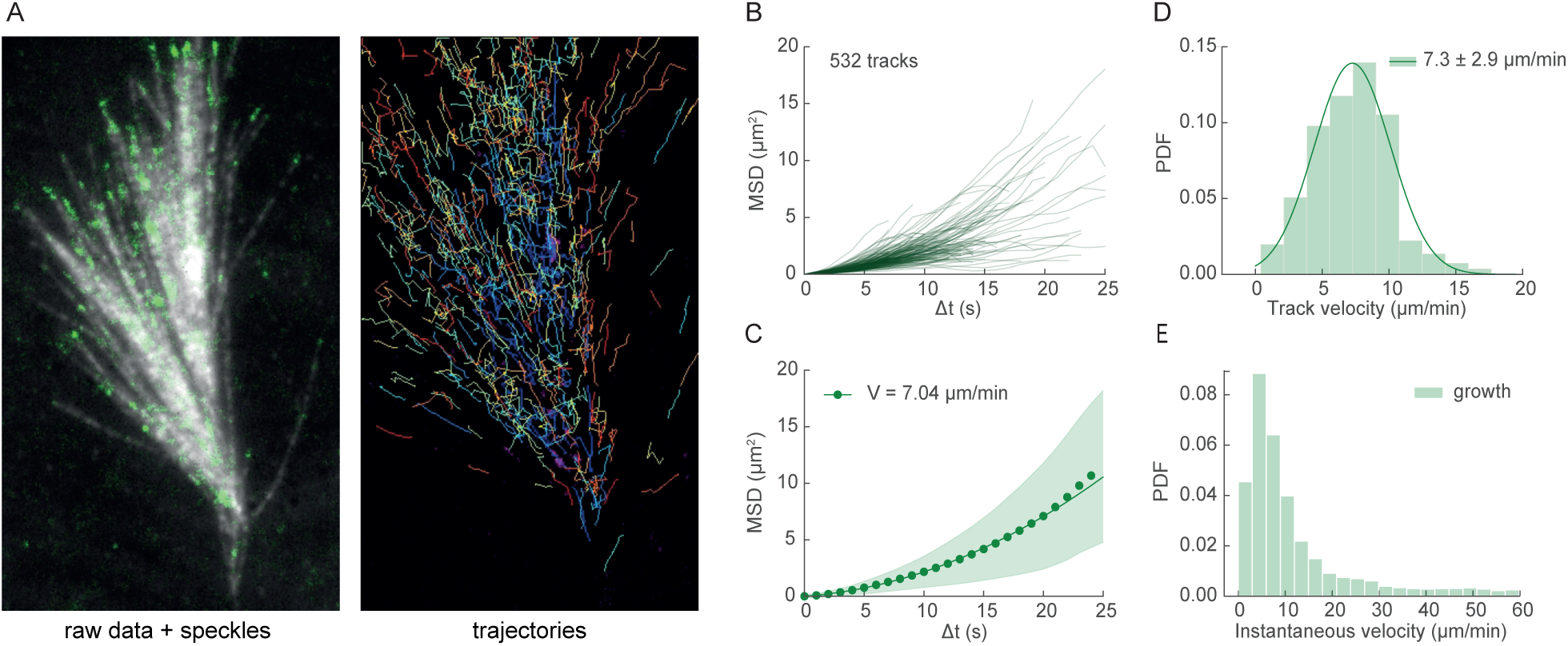
Quantitative analysis of microtubule growth. **(A)** Snapshot of a microtubule branching time-lapse experiment merged with fluorescent speckles, corresponding to the growing ends (left) and examples for trajectories obtained after performing automated tracking (right). Scale bar is 2µm. **(B-E)** Summary of the analysis output **(B)** MSD curves for individual trajectories and **(C)** the weighted mean MSD curve for growing microtubule ends, where the solid line represents a fit of a quadratic equation (eq.3) to the data, to estimate the mean velocity. Shaded bars represent the standard deviation of all tracks. **(D)** A Gaussian function (solid line) is fitted to the distribution of all MSD to estimate the mean velocity value. **(E)** Distribution of velocity values obtained from the step size displacements of speckles between consecutive time points.

## V. Conclusions

We present a framework for the computational analysis of treadmilling filaments, in particular the rate of polymerization and depolymerization of membrane-bound FtsZ filaments. Although we used this approach to quantitatively characterize growth and shrinkage of treadmilling FtsZ filaments *in vitro*, this approach is applicable to study the polymerization dynamics of other cytoskeletal systems as well. In contrast to previous methods, this method can be applied on any time-lapse movie of homogeneously labelled filaments as it does not require specific markers for polymer ends. It also allows to quantify both, growing and shrinking rates of dynamic cytoskeletal filaments. Furthermore, it can be easily extended with more complex analyses, for example to generate spatiotemporal maps of filament dynamics in living cells. In fact, combined with an appropriate detection procedure, the approach used here should also allow to visualize and track motion on every spatial and temporal scale, not only cytoskeletal structures *in vivo* and *in vitro*, but also crawling cells and migrating animals.

In sum, the method described here surpasses the limitations of kymographs, as it allows for a non-biased quantification of dynamic processes in an automated fashion.

## Acknowledgements

We kindly thank Franziska Decker, Benjamin Dalton and Jan Brugués (Max Planck Institute, Dresden, Germany) for providing us with a time lapse of movie of microtubules and also for their feedback on our manuscript. We thank all Loose lab members for support and useful discussions, in particular to Natalia Baranova for the critical review of our methodology in general. This work was supported by a European Research Council (ERC) grant awarded to Martin Loose (ERC-2015-StG-679239) and a Boehringer Ingelheim Fonds (BIF) PhD fellowship awarded to Paulo Caldas.

## Author contributions

<u>Project Management</u>: M.L; <u>Conceptualization and Design</u>: M.L, P.C and P.R; <u>Experimental Data</u> <u>Collection and Image analysis</u>: P.C and P.R; <u>ImageJ and Python Programming</u>: C.S and P.C; <u>Data interpretation</u>: M.L, P.C and PR; <u>Data Visualization</u>: P.C, P.R and C.S; <u>Manuscript</u> <u>Drafting</u>: P.C and M.L; <u>Critical Revision of the Manuscript</u>: All authors; Final version was read by all authors and approved to be published.

## Competing interests

The authors declare no competing interests.

